# Mitochondrial respiratory capacity in kidney podocytes is high, age-dependent, and sexually dimorphic

**DOI:** 10.1101/2024.11.24.625104

**Authors:** Matthew D. Campbell, Monica Sanchez-Contreras, Britta D. Sibley, Phoebe Keiser, Carla Ruiz Sanchez, Carolyn N. Mann, David J. Marcinek, Behzad Najafian, Mariya T. Sweetwyne

## Abstract

Whether and how podocytes depend on mitochondria across their long post-mitotic lifespan is yet unclear. With limited cell numbers and broad kidney distribution, isolation of podocyte mitochondria typically requires first isolating podocytes themselves. Disassociation of podocytes from their basement membrane, however, recapitulates an injured state that may stress mitochondria. To address this, we crossed floxed hemagglutinin (HA) -mitochondria tagged (MITO-Tag) mice with those expressing Cre in either podocytes (NPHS2) or distal tubule and collecting duct (CDH16), thus allowing for rapid, kidney cell-specific, isolation of mitochondria via immunoprecipitation. Mitochondrial respiration in fresh isolates from young (4-7 mo) and aged (22-26 mo) mice of both sexes demonstrated several previously unreported significant differences between podocyte and tubule mitochondria. First, although podocytes contain fewer mitochondria than do tubule cells, mitochondria isolated from podocytes averaged twice the respiratory capacity of tubule mitochondria when normalized to mitochondrial content by citrate synthase (CS) levels. Second, age-related decline in respiration was detected only in podocyte mitochondria and only in aged male mice. Finally, disassociating podocytes for cell culture initiates functional decline in mitochondria as those from cultured primary podocytes have half the respiratory capacity, but twice the hydrogen peroxide production of podocyte mitochondria isolated directly from fresh kidneys. Thus, podocytes maintain sexually dimorphic mitochondria with greater oxidative phosphorylation capacity than mitochondria-dependent tubules per organelle. Previous studies may not have detected these differences due to reliance on podocyte cell culture conditions, which results in artifactual suppression of mitochondrial function.

## INTRODUCTION

Kidneys are among the most mitochondria rich and energetically demanding organs due primarily to their high resting metabolic rate. As such, dysregulation of cellular metabolism through mitochondrial dysfunction is a hallmark of acute kidney injury^1,2^, chronic kidney diseases^3,4^, and kidney aging^5,6^. The structural and functional changes to mitochondria that underlie dysfunction likely differ based on the physiology of each of the more than 30 unique cell types in the kidney. Most studied in this context are the mitochondria rich, and mitochondrially dependent, kidney tubule epithelial cells, which are responsible for the active transport driving reabsorption and secretion of the nephron. Tubules comprise about 90% of the kidney cortex by volume and are ranked in the top three most mitochondrially endowed cells of the body, depending on both oxidative phosphorylation and fatty acid oxidation for their high demand of ATP generation. This endowment and dependance has driven our recognition of mitochondria as critical for kidney function, but mitochondria studies that relied on whole kidney extracts essentially only capture the propensity of tubule cells in the kidney, and of mitochondria in the tubule cells. This tubule-centric view of kidney mitochondria has confounded our understanding of mitochondrial dependance in less abundant unique kidney epithelial cell types, such as the podocyte.

Podocytes, unlike tubule cells, are post-mitotic epithelia that are essential for formation of the glomerular filtration slit but do not perform active solute transport across the nephron. Multiple studies in mice have shown that mitochondria regulate detrimental changes to podocytes with disease, age or injury ^5,7–9^. Our own work with the mitochondrially-targeted peptide, Elamipretide (previously SS-31 and Bendavia), shows that 8 weeks of systemic treatment prevents progression of glomerulosclerosis and significantly lowered markers of podocyte injury, glomerular Desmin protein expression and podocyte hypertrophy, while increasing expression and glomerular distribution of the podocyte cytoskeletal protein, synaptopodin^5^. In humans, structural disruption to podocyte mitochondria is detected in diabetic patients^7^ and specific inherited mutations in mitochondrial DNA can cause podocytopathy.

Despite this, evidence for mitochondrial dependance of healthy podocytes is sparse. Uninjured podocytes in young mice have thus far have shown little dependence on mitochondrial energetics, appearing to rely primarily on glycolysis for ATP production. This, coupled with relatively low podocyte mitochondrial endowment, has led to the paradigm that mitochondria are non-essential for podocyte health ^11,12^. The current understanding of how podocytes specifically rely on mitochondria has been largely shaped by *in vitro* experiments with primary and immortalized cell lines or single target knock-out animals. The high propensity of podocyte mitochondrial damage in glomerular diseases led us to hypothesize that mitochondrial regulation of cell functions beyond oxidative phosphorylation might explain why mitochondria intervention is protective of aged and diabetic podocytes. Furthermore, we suspected that the cellular disassociation and culture required to isolate podocyte mitochondria away from the ubiquitous tubule mitochondria, could be inducing metabolic changes and other physiologically relevant mechanisms in podocyte mitochondria.

Here we explored the functional roles of mitochondria in podocyte health as was not previously possible due to difficulty in accessing podocyte mitochondria *in vivo* and in analyzing them within the background of overwhelming mitochondrial density and activity from kidney tubules. To do this, we took advantage of the MITO-Tag mouse model, which we crossed with mice expressing Cre in either the podocyte or Cdh16/Ksp1 expressing tubule cells, thus generating new mouse lines that allow us to rapidly isolate podocyte or tubule mitochondria without first disassociating the kidney into single cells. Our intention was to use these mice in injury models to determine how mitochondria in different kidney cell types respond to insult. In confirming the expression patterns of our mice and the efficacy of our mitochondrial isolation protocol, however, we found a surprising difference in the fundamental respiratory capacity and sex-specific aging decline of mitochondria from podocytes, relative to kidney tubule mitochondria.

## Methods

### Urinalysis

Urines were collected from all mice +/- one week from the corresponding age group. Albumin concentrations were measured using the Albuwell M ELISA Kit (Ethos Biosciences, Logan Township, NJ), and Creatinine concentration was measured using the Creatinine Colorimetric Assay Kit (Cayman Chemical, Ann Arbor, MI). For all initial analyses of samples from young mice, urines were diluted to 1:21 for albumin analysis, or to 1:11 for creatinine analysis, and then analyzed according to manufacturer instructions. For all initial analyses of samples from old mice, urines were diluted to 1:35 for albumin analysis and 1:11 for creatinine analysis. Samples exceeding a CV of 10% for replicates were re-run and excluded if the CV remained >10%.

### Animal Care

All mice used in this study are on the C57BL/6j background. The NPHS2-cre transgene was introduced into the colony with mice of the B6.Cg-Tg(NPHS2-cre)295Lbh/J strain (Jackson Laboratory, Bar Harbor, ME), the CDH16-cre transgene with mice of the B6.Cg-Tg(Cdh16-cre)91Igr/J strain (Jackson Laboratory), the MITO-GFP mutation with mice of the B6N.Cg-Gt(ROSA)26Sortm1(CAG-EGFP*)Thm/J strain (Jackson Laboratory), and the MITO-HA mutation with mice of the B6.Cg-Gt(ROSA)26Sortm1(CAG-EGFP)Brsy/J strain (Jackson Laboratory). Mice were housed at 20°C under diurnal conditions with *ad libitum* food and water in an AAALAC-accredited facility with supervision by the Institutional Animal Care and Use Committee. Genotypes of all animals in this study were obtained through the automated genotyping service provided by TransnetYX.

### Mitochondrial Isolation and Enrichment from Whole Kidney Tissue

Mice were euthanized by cervical dislocation and their kidneys immediately collected, and decapsulated. 200 mg of kidney was cut into ∼5 mm pieces and transferred to a pre-chilled Dounce homogenizer with 1 mL of 1x KPBS (136 mM KCl, 10 mM KH2PO4, pH 7.25 in MilliQ water). Tissue was homogenized for 2 min on ice. Homogenate was filtered through a 70 µm cell strainer, diluted in 3 mL of 1x KPBS and centrifuged for 2 min at 1,000 x g at 4°C. To isolate cell specific kidney mitochondria and total kidney mitochondria from the same homogenate, supernatant from each mouse was split into two equal volumes brought to 10 mL with Miltenyi 1x Separation Buffer (Miltenyi, Bergisch Gladbach, Germany). Suspensions were immunopreciptated with either 50 μL of anti-HA magnetic microbeads (µMACS HA Isolation Kit, Miltenyi) or 50 μL anti-TOMM22 magnetic microbeads (Mouse Mitochondria Isolation Kit, Miltenyi) for 45 min at 4°C under continuous rotation. Immunoprecipitants were loaded into an LS column (Miltenyi), washed and eluted in Separation Buffer as directed the manufacturer. Separation Buffer was removed by centrifuging at 13,000×g for 2 minutes at 4°C, discarding the supernatant and resuspending in Respiration Buffer (0.5 mM EGTA, 20 mM taurine, 3 mM MgCl2, 110 mM Sucrose, 60 mM K-MES, 20 mM HEPES, 10 mM KH2PO4, 1mg/mL BSA, pH 7.1). The wash step was repeated, and mitochondria pellets were resuspended in 100 μL of Respiration Buffer to be used immediately for mitochondrial function analysis or to be stored at - 80°C for non-functional assays.

### Immunofluorescence

Frozen OCT embedded kidneys were sectioned and fixed in 4% paraformaldehyde for 10 minutes. Fixed slides were rinsed twice in PBS and permeabilized with 50 μL of Triton X-100 (Sigma, St. Louis, Missouri) for 8 minutes. After permeabilization, slides were rinsed twice with PBS before adding blocking buffer (2% Goat serum, 1% BSA) for 15 minutes. Goat anti-mouse HA primary antibody (Novus, Centennial, Colorado) was diluted at 1:750 in dilution buffer (1% BSA, 0.05% Tween-20, 1X TBS) and incubated at 4℃ overnight. Slides were rinsed twice with PBS + 0.05% Tween-20 (Sigma) and incubated in 1:200 Donkey anti-Goat Alexa Fluor 488 secondary antibody (Invitrogen, Carlsbad, California) for 1 hour. Slides were rinsed twice with PBS and 1mM Hoechst solution (ThermoFisher, Waltham, MA) was applied for 5 minutes before final rinse in PBS and mounting with Vectashield (Vector Laboratories, Burlingame, California).

### Periodic acid-Schiff stain

Mouse renal biopsies were fixed in 4% paraformaldehyde at 4℃ overnight followed by 70% ethanol and embedded in paraffin. 4 µm sections were stained with periodic acid-Schiff and hematoxylin by the Pathology Research Services laboratory at UW. Sections were blinded and imaged using an EVOS FL Auto 2. 40X images of glomeruli were taken at random points of the cortex for all sections. An image that was representative of the morphology of each section was selected to be included in figures.

### Transmission Electron Microscopy

Mouse kidney tissue was processed per the Ellisman protocol, (dx.doi.org/10.17504/protocols.io.36wgq7je5vk5/v2). Thick 1 µm sections were cut using an RMC ultramicrotome, stained with toluidine blue for morphology evaluation, and thin 80 nm sections were imaged under a JEOL 1011 TEM at various magnifications for further analysis.

### Respirometry/Fluorometry

Following mitochondrial isolation and enrichment, 45-50ug of purified mitochondria was added to a 2mL chamber of an Oxygraph 2-K dual respirometer/fluorometer (Oroboros Instruments, Innsbruck, AT) in respiration buffer (0.5 mM EGTA, 20 mM taurine, 3 mM MgCl_2_, 110 mM Sucrose, 60 mM K-MES, 20 mM Hepes, 10 mM KH2PO4, 1mg/mL BSA, pH 7.1) at 37°C, with 750 rpm stirring. O_2_ concentration was maintained between 50 and 250 µM. Respiratory states were measured with the following sequential titrations of substrates and inhibitors (final concentrations): malate (0.1 mM), palmitoyl carnitine (70 µM), sub-saturating ADP (50 µM), saturating ADP (2.5 mM), malate (1 mM) and pyruvate (5 mM), glutamate (10 mM), cytochrome C (2.5 µM), succinate (2.5 mM), succinate (7.5 mM), rotenone (0.5 µM), antimycin A (2.5 µM), Ascorbate (2 mM) and TMPD (0.5mM), Sodium Azide (100 mM). All data was analyzed using Datlab 7.4.0.4 (Oroboros Instruments) and GraphPad Prism 10.2.2. Respirometry values were compared using one-way ANOVA, with a Tukey’s multiple comparisons test.

### Western blot

Mitochondrial Isolation Blots: Whole tissue lysates were prepared from 20mg of flash-frozen kidney cortex tissue using Qiagen TissueLyser at 4°C. Whole tissue powder and mitochondrial isolations were lysed in RIPA buffer with protease/phosphatase inhibitor cocktail (ThermoFisher, Waltham, MA). Protein concentrations were determined by BCA (ThermoFisher). 15 µg of each lysate was mixed with 1x Laemmli buffer (ThermoFisher) and 1x Reducing Agent (ThermoFisher) and brought up to 25 μL with RIPA buffer before boiling at 95°C for 10 min. Samples were loaded onto 4-12% Bis-Tris NuPAGE gels (ThermoFisher) and transferred to 0.45 µM PVDF membranes (MilliporeSigma, Burlington, MA). Total loaded protein was determined via Ponceau S staining. Membranes were blotted sequentially, and stripping was avoided where possible to conserve protein samples. Blots were imaged using the iBright FL1500 and images were analyzed via ImageJ (National Institutes of Health, Bethesda, MD, USA). All antibodies used are listed in supplementary methods

### Glomerular isolation and culture

Kidneys were minced and digested for 20 minutes in 1 mg/mL Collagenase A (Worthington Biochemical, Lakewood, New Jersey). Isolates were pushed through 50 mL conical tube sterile 100 µm cell strainers and washed with cold 35 mL HBSS (Gibco, Grand Island, New York) + Ca^2+^, Mg^2+^ and 1 mM HEPES. Flow through was centrifuged for 4 minutes at 2,000 rpm in a 4°C swinging bucket rotor. Supernatant was discarded and red blood cells were lysed in 4 mL of RBC lysis buffer (Invitrogen, Carlsbad, California) for 10 minutes on ice. Lysis was quenched with 40 mL of cold HBSS, and resuspended kidney slurry was poured through a 70 μm pore size cell strainer. Glomeruli in flow-throughs were collected on a 40 μm cell strainer and washed 2x in 25 mL of cold HBSS to remove excess digest buffers and cell debris. Glomeruli were cultured in equal number on a 60 mm tissue culture treated dish at 37°C, 5% CO_2_ and 5% O_2_ in RPMI 1640 (Gibco) with 15% FBS (Gibco) and penicillin streptomycin (Corning, Manassas, Virginia) for 72 hours. Day 3, cultures were washed with sterile PBS (Corning) to remove unattached cells and debris and FBS lowered to 10%. On day 7, cells were washed twice in PBS and collected with a rubber scraper into 1.5 mL of MACS Lysis Buffer provided with the mitochondria isolation kit (Miltenyi, Germany). Cell lysates were passed 5x through a 25-gauge needle to shear cells. Lysates were resuspended in 4mL of KPBS, and mitochondria were then isolated exactly as described for whole kidney tissue starting from KPBS suspension after homogenization.

## RESULTS

### Characterization of Podocin (NPHS2)-MitoTAG and Tubule (CDH16)-MitoTAG mice

The MITO-Tag mice, B6.Cg-*Gt(ROSA)26Sor^tm1(CAG-EGFP)Brsy^*/J ^21^ (hereafter referred to as **mtHA)**, were each crossed with either B6.Cg-Tg(NPHS2-cre)295Lbh/J (**NPHS2**) or B6.Cg-Tg(Cdh16-cre)91Igr/J (**CDH16**) for expression in the podocyte ^22^ or mixed tubule ^23^ cells respectively. Heterozygosity for both the MITO-Tag and the Cre is sufficient for tissue level expression which was confirmed by observing appropriate cell localization to either the podocyte **(Fig 1)** or loop of Henle, distal and collecting duct cells **(distal tubule shown: Fig 1**) by immunofluorescence (IF) against the HA-tag. Having successfully generated kidney mtHA mouse lines, we also introduced a MITO-Tag enhanced green fluorescent protein (EGFP)^29^ tag on the outer membrane of mitochondria in either the podocyte or tubule cells of the kidney by crossing the Cre lines with a commercially available mouse, B6N.Cg-*Gt(ROSA)26Sor^*tm1(CAG-EGFP*)Thm*^*/J ^29^ (**mtGFP)**. Our intention was to use the mtHA kidney mouse lines for mitochondrial pulldown **(Fig. 1)** and the mtGFP mouse lines to perform live tissue imaging of mitochondria dynamics. Of note, the two different MITO-Tag strains were generated by separate research groups, but both used identical strategies to target their tags to the outer mitochondrial membrane via transgenic expression of a fusion protein that includes the same transmembrane domain (aa109-145) of the rat Omp25/Synnj2bp mitochondrial protein to target the mitochondrial outer membrane^10,11^. Although both groups generating the MITO-Tag mice included an EGFP sequence in their constructs, only the mtGFP group (Misgeld) succeeded in deriving a construct that encodes readily detectable EGFP fluorescence in unfixed tissue^10,12^. We confirmed that crossing their mouse with the kidney Cre lines resulted in appropriate kidney cell-specific and robust fluorescence of the EGFP-tag in both the NPHS2^Cre^- and CDH16^Cre^- mtGFP crosses (**Fig. 1**).

**Figure 1.**
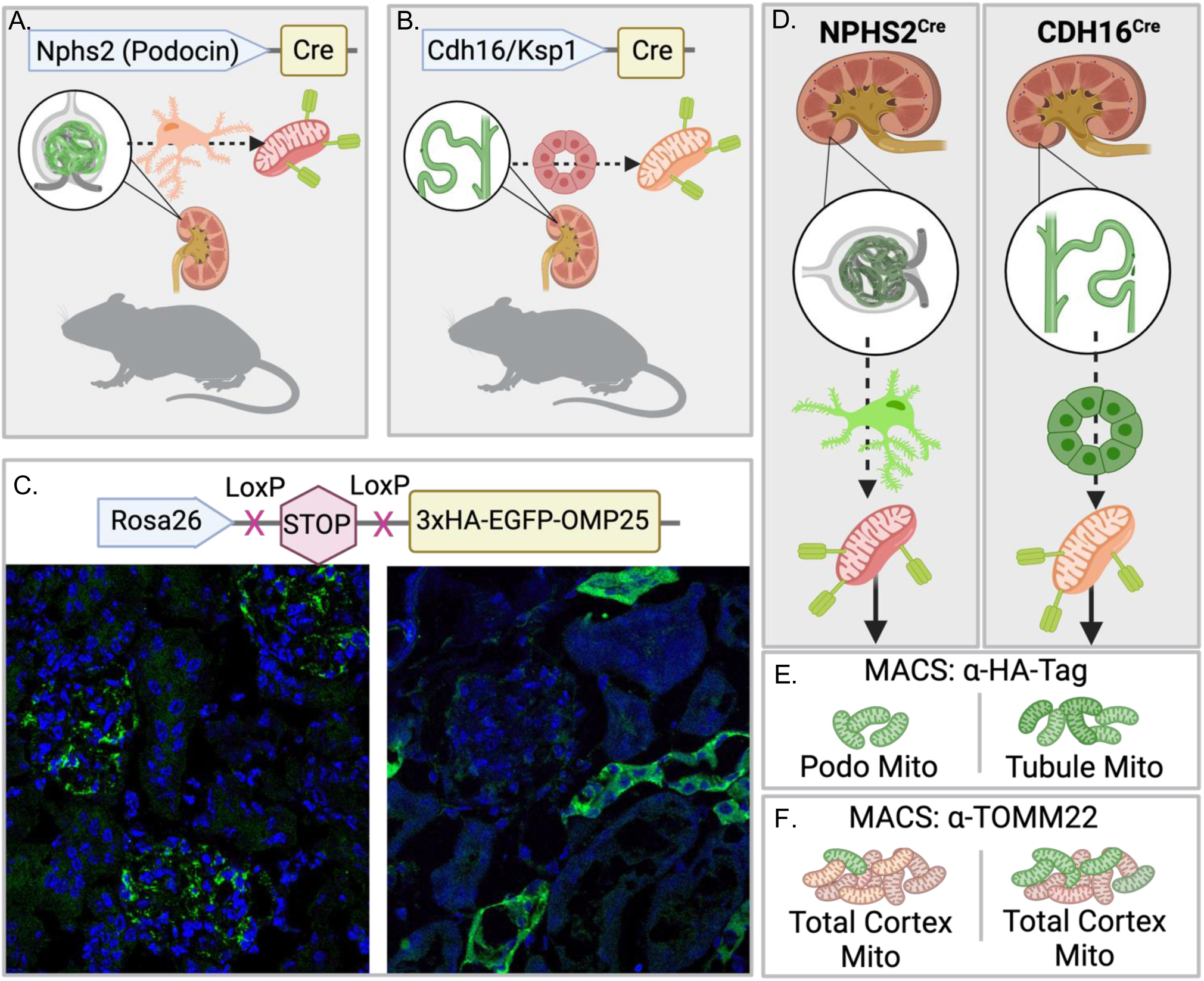
Breeding scheme and characterization of Podocin NPHS2-mtHA and Tubule CDH16-mtHA mice. Expression of tagged mitochondria in podocytes or tubule cells was obtained by crossing strains expressing Cre recombinase directed by either **(A)** the human podocin promoter/enhancer to podocytes (Nphs2-Cre) or **(B)** by mouse cadherin 16 promoter to tubule cells (Cdh16-Cre). **(C)** Crossing Nphs2-Cre and Cdh16-Cre cell lines to MITO-Tag-HA mice carrying the Gt(ROSA)26Sor^tm1(CAG-EGFP)Brsy^ construct where 3xHA-EGFP-OMP25 is the “mtHA” cassette resulted in correct expression localization, which was confirmed by immunofluorescence against HA-tag in either podocytes **(C, left)** or distal/convoluted tubules **(C, right)**. **(D)** Schematic of rapid cell-specific mitochondrial isolation using Miltenyi MACS immunoprecipitation of either HA-tagged mitochondria (cell-specific, **E**) or TOMM22 for indiscriminate pulldown of all kidney cortex mitochondria (**F**).

To rapidly isolate mitochondria from either podocytes or tubule cells we modified a protocol using <u>m</u>agnetic <u>a</u>ctivated <u>c</u>ell-<u>s</u>orting (MACS, Miltenyi Biotec) columns, and two different isolation kits, for pull-down of mitochondria against the HA-tag **(Fig 1)**. As a control, we also isolated total mitochondria from a portion of the same kidney cortices via an indiscriminate mitochondria MACS pull-down against outer membrane mitochondrial protein, TOMM22 **(Fig 1)**.

### Functional EGFP MitoTAG, but not HA-MitoTAG, causes renal failure through podocyte toxicity

Most *in vivo* studies of kidney function use experimental mice once they reach sexual maturity soon after 4-7 weeks of age, which is still quite juvenile developmentally.^13,14^ Because we study aging in the adult kidney, we age our mice to at least 16-weeks-old, analyzing them in the young adult group between 4-7 months of age. While generating and aging this new colony, we observed that some of the NPHS2-mtGFP mice were dying at young ages starting as early as 2 months old. This was unexpected as we had used one male and two female breeder mice from each of the four founding lines to generate our colony. To make our genotype crosses, we had arranged breeding trios so that each MITO-Tag male was mated with a NPHS2^Cre^ and a CDH16^Cre^ female, and then rotated the breeders so that each MITO-Tag female birthed at least one litter from each Cre line as well. This breeding strategy was intentional to maximize the relatedness between our four different crossed lines where possible, and it meant that we were certain that the expression of the mtGFP MITO-Tag itself was not inherently toxic since we did not observe any early deaths in the CHD16-mtGFP crosses, which were offspring of the same mtGFP founding sire or dam as for the affected NPHS2-mtGFP mice.

A survival curve of cohorts intended for late age studies showed that survival of the NPHS2-GFP mice was only 50% by 8 months of age (9 of 18 mice) **(Fig 2A)**. In comparison, the same attrition was not observed in controls (97% survival, 58 of 60), nor in the NPHS2-mtHA (99% survival, 108 of 109), CDH16-mtHA (100%, n=138) or the CDH16-mtGFP mice (100%, n=16) (**Fig 2A**). We continued to follow the survival of our control and mtHA-TAG mouse lines to 24-months-old, which is a standard late-age time point for our aging studies. There was no significant difference in the survival of either NPHS2-mtHA (66% survival, 25 of 38) or CDH16-mtHA mice (survival, 78%, 18 of 23) relative to control animals (71% survival, 12 of 14) **(Fig. 2B)**.

**Figure 2.**
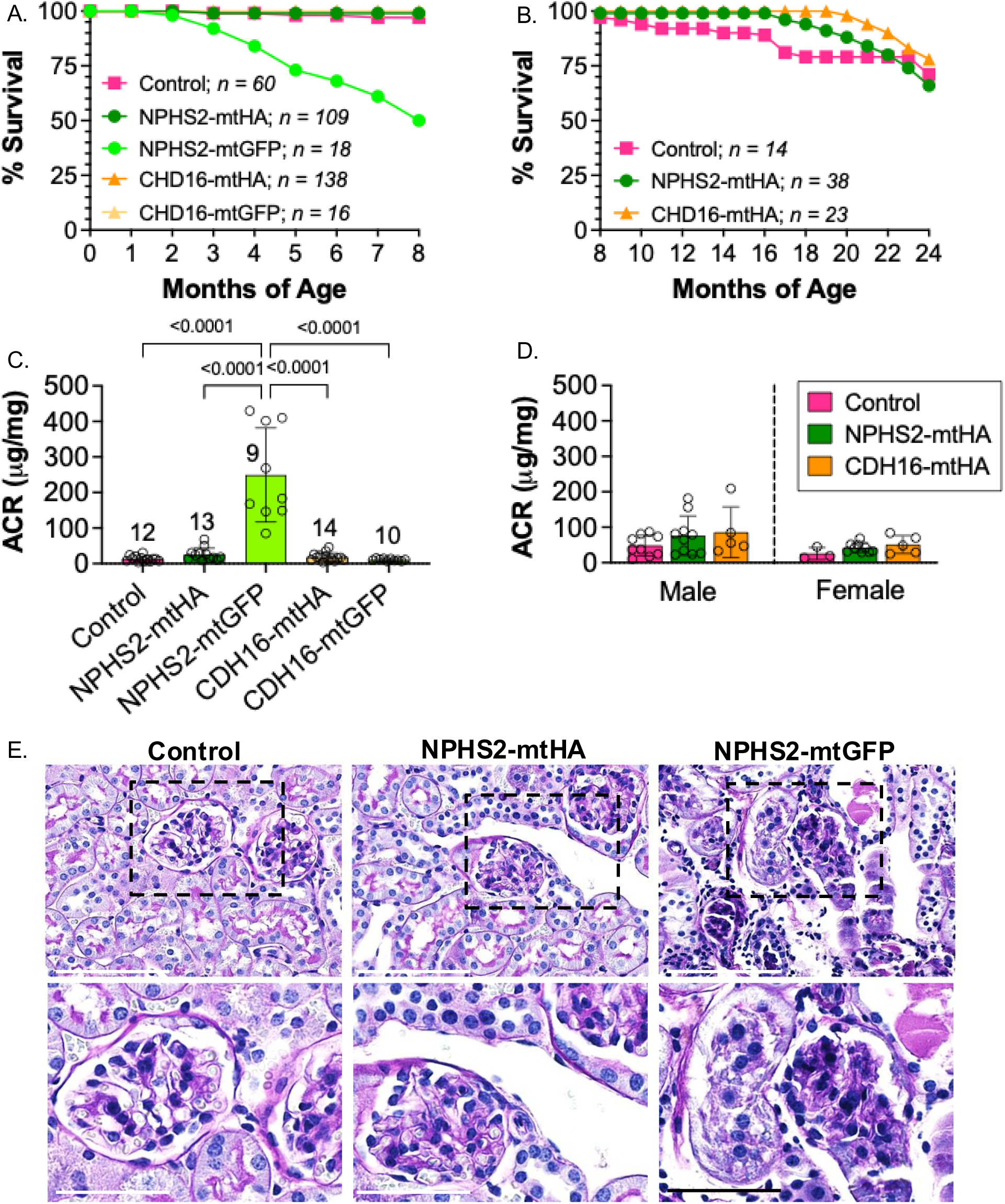
MITO-Tag-GFP tag is detrimental when expressed in the podocyte (NPHS2-mtGFP) but not tubule (CDH16-mtGFP) mice. **A.** Survival curve of Podocin NPHS2-mtHA and Tubule CDH16-mtHA compared to NPHS2-mtGFP and Tubule CDH16-mtGFP shows increased mortality in the NPHS2-mtGFP strain before 8 mo. **B.** Survival of NPHS2-mtHA and CDH16-mtHA strains used for aging studies are comparable to control mice that do not express a MITO-Tag. **C.** Urinary Albumin/Creatine ratio (ACR) is significantly elevated in NPHS2-mtGFP strain indicating proteinuria resulting from glomerular filtration deficiency. **D.** No ACR differences from control are detected in 20–24-month-old NPHS2-mtHA or CDH16-mtHA mice, indicating that glomerular filtration is preserved in mtHA males and females into late age. **E.** Periodic-acid Schiff stain of 5-month-old male mouse kidneys shows normal tubule and glomerular architecture in control and NPHS2-mtHA mice. NPHS2-mtGFP mice show severe structural abnormalities of the kidney characterized by collapsing glomeruli and atrophy of surrounding tubules. Top panel scale bar = 100 µm and bottom panel scale bar = 50 µm. Error bars = +/- SD, significance determined as <0.05 by one-way ANOVA.

The long-term survival of our mtHA mice suggests no overt disruption to the kidney with the mtHA tag expression, but we also confirmed that there were no functional anomalies by checking for proteinuria. Urinary albumin to creatinine ratio (ACR) was measured from mice 4-8-months-old in all kidney MITO-Tag mouse lines and controls. Only in NPHS2-mtGFP mice was the ACR elevated (mean of 249 µg/mg) relative to control (mean = 13.38 µg/mg), NPHS2-mtHA (mean = 25.71 µg/mg), CHD16-mtHA (mean = 18.15 µg/mg), and CHD16-mtGFP mice (10.65 µg/mg) (**Fig 2C**). Two NPHS2-mtGFP mice were sacrificed for rapid necropsy by University of Washington Veterinary Services with the primary finding being severe diffuse glomerulonephritis with tubular atrophy. We stained mouse kidney biopsies from 5-month-old male mice with periodic-acid Schiff stain (PAS) and found that both control and NPHS2-mtHA mice had grossly normal kidneys with thin and distinct basement membranes, healthy glomeruli with clear capillary loops and no mesangial expansion (**Fig 2E**). On the other hand, the NPHS2-mtGFP mice had compacted glomeruli and tubular atrophy with casts as also reported by necropsy **(Fig 2E)**.

3D-structured illumination microscopy (3D-SIM) analysis of the podocyte filtration slit density (NIPOKA, Germany) showed podocyte injury in NPHS2-mtGFP mice with disrupted filtration slit formation (**Fig 3A-C**). Transmission electron microscopy of glomeruli in the NPHS2-mtGFP mice showed that podocytes had thickened foot processes and an unusual flower-like aggregation of mitochondria (**Fig 3D, E**), whereas TEM in NPHS2-mtHA mice revealed normal size and shape foot processes and podocyte mitochondria (**Fig 3F,G**). Based on these collective results **(Figs 2 & 3)**, we concluded that the mtHA tag was not disruptive to mitochondria or cell function in either podocytes or tubule cells, however, we chose not to continue analysis with the mtGFP mouse lines. Despite this, the mtGFP mouse lines were useful in demonstrating that podocytes are particularly sensitive to mitochondrial disruption to a degree that can cause total renal failure and death if the dysfunction is robust. We continued our studies of podocyte versus tubule mitochondria using mtHA kidney mouse lines with the MACS pulldown method to analyze the functional respiratory capacity kidney cell-specific mitochondria.

**Figure 3.**
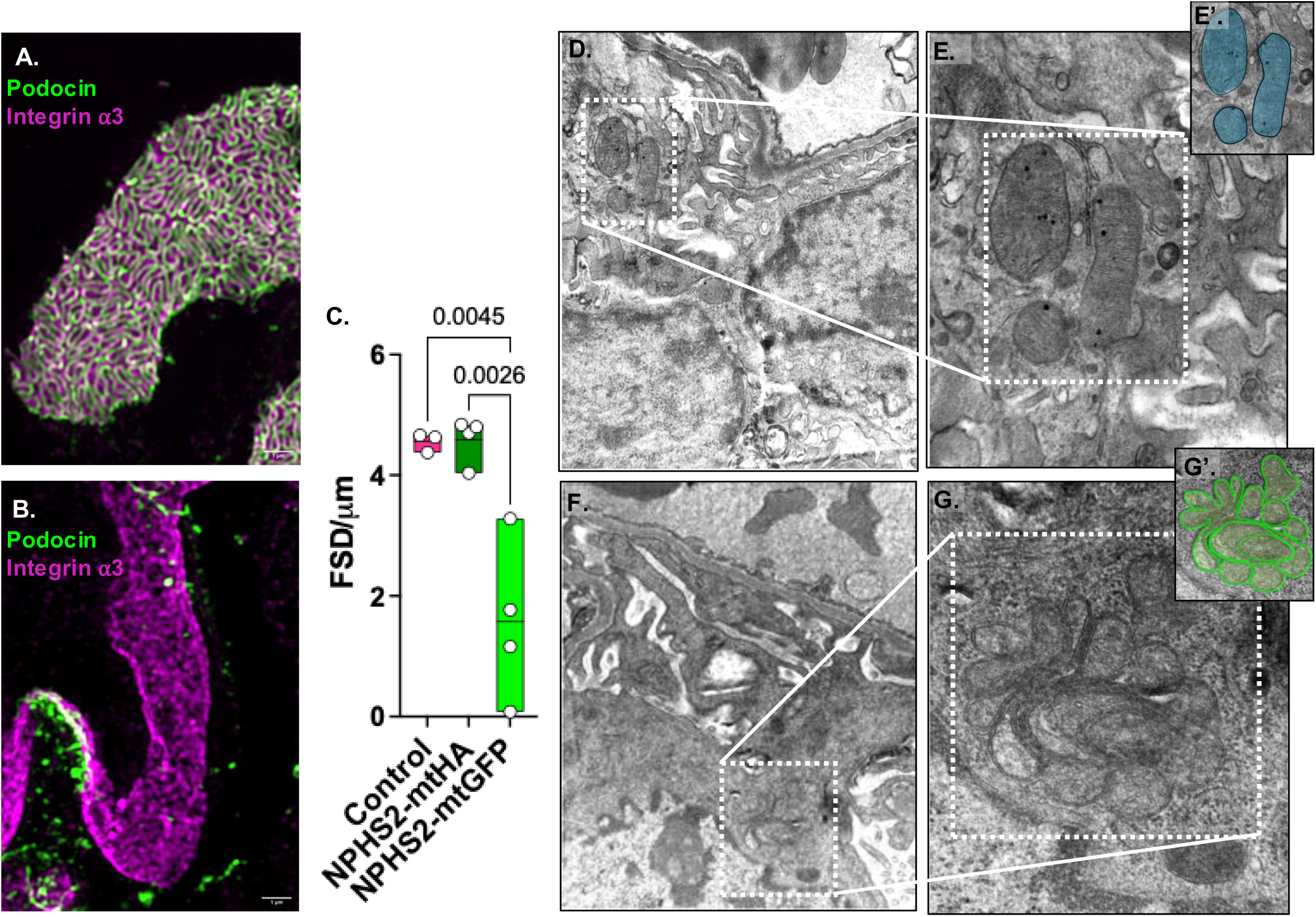
Ultra-structural abnormalities of podocyte feet processes and their mitochondria by EM suggestive of podocyte injury in NPHS2-mtGFP mice. **(A)** Green = Podocin; Magenta = Integrin α3, Super resolution structured illumination microscopy (SIM) example of NPHS2-mtHA foot process with normal podocyte filtration slit diaphragm morphology. **(B)** loss of podocin from the podocyte slit diaphragm is seen in NPHS2-mtGFP mice. **(C)** Filtration slit density (FSD) is severely reduced in NPHS2-mtGFP mice as measured by the average FSD of 20 randomly selected glomeruli per mouse. **(D – E’) Podocyte mitochondria from NPHS2-mtHA have normal morphology by transmission electron microscopy. (F-G’)** NPHS2-mtGFP podocyte foot processes are and have abnormally large mitochondria clusters lacking almost all cristae. **(E’ & G’)** show color overlay to assist with identifying mitochondria in micrographs.

### Assessment of mitochondrial pull-down from HA-tag

The initial report using the MITO-Tag mice in liver showed that the HA-tag isolation method produced functional mitochondria with some co-elution of other cell organelles known to associate closely with mitochondria such as endoplasmic reticulum, peroxisomes and lysosomes^11^. To assess the composition of our mitochondria MACS pulldowns, we performed immunoblotting of HA (cell-specific mitochondria) and TOMM22 (total kidney mitochondria) pull-downs, as well as whole kidney homogenate from adult male mice (11-12 mo). We also included control mice that did not express a HA-tag with either TOMM22 pull-down or whole kidney homogenate. First, we stained the blots with Ponceau S for relative assessment of total protein. Although all wells were loaded for equal protein concentration based on BCA assay, variation in total protein was detected across lanes **(Fig 4A)**. We next blotted using antibodies against mitochondrial proteins TOMM20 and ATP5a. Expression of mitochondrial proteins, TOMM20 and ATP5a were detected across all samples as expected, but the two proteins were not in congruence with respect to the relative concentrations between samples when normalized to total protein. Because TOMM20 concentration appeared to more closely follow total protein expression we chose to use TOMM20 densitometry to normalize the remaining immunoblot targets against mitochondrial content.

**Figure 4.**
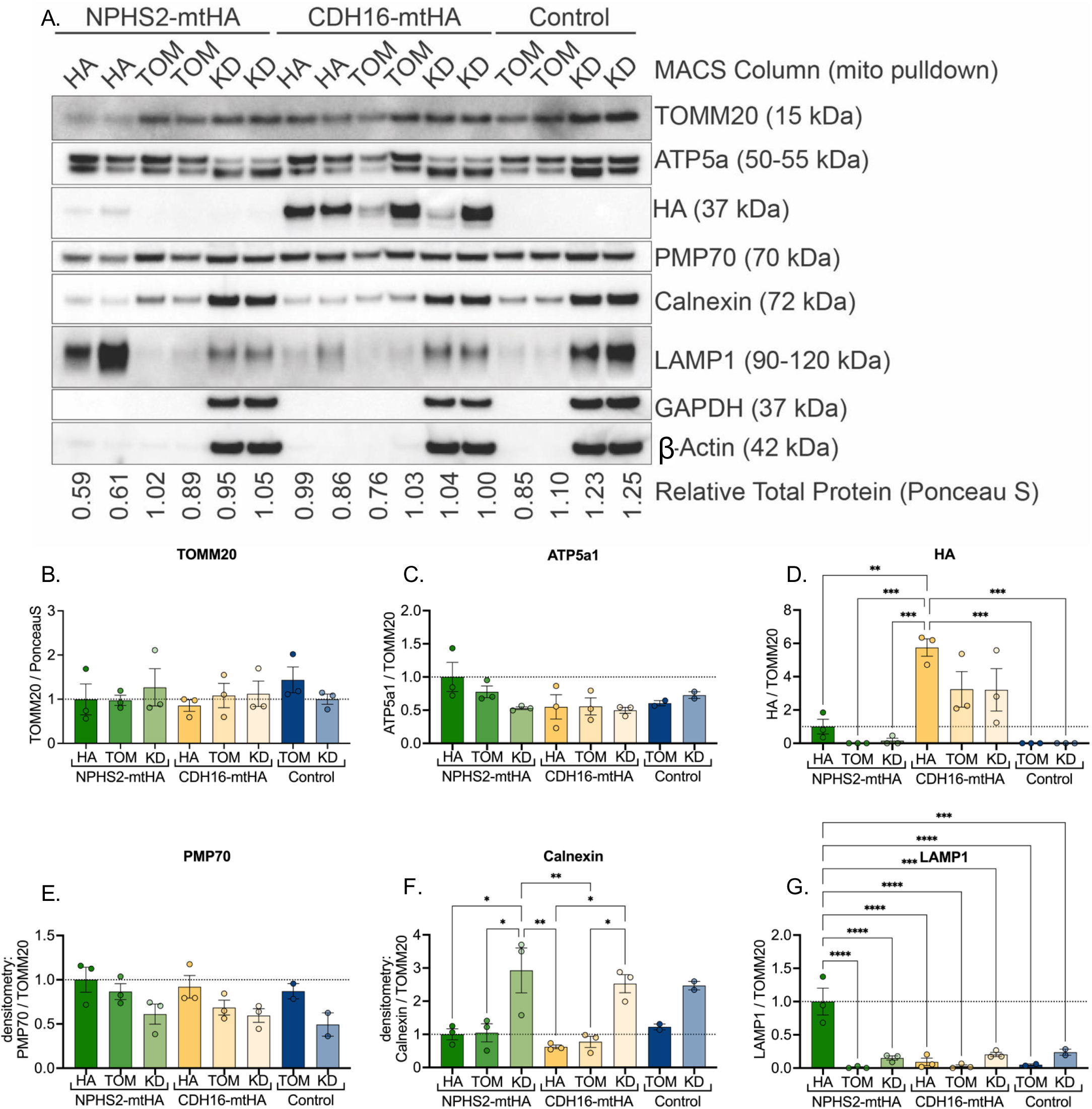
Assessment of the selective isolation of mitochondria using MACS immunolabeling. **(A)** Western blots of protein isolates obtained from the MACS immunolabeling and magnetic separation with anti-HA (HA) or anti-TOMM22 microbeads (TOM), or whole kidney tissue homogenates (KD). Target proteins: TOMM20 and ATP5a1 (mitochondria), HA (HA-tag), PMP70 (peroxisomal membrane), Calnexin (endoplasmic reticulum), LAMP1 (lysosomes), GAPDH (cytoplasm), β-Actin (cytoplasm). **(B to G)** Quantification by densitometry of the western blot intensity. In all blots the average densitometry of NPHS2-mtHA HA pulldown is set to baseline 1 and all other group are expressed as relative values. **(B)** TOMM20 protein densitometry is normalized to total protein from Ponceau S stain (values in A). (C-G) All blots normalized to TOMM20 densitometry. Error bars = +/- SD, statistical significance determined as <0.05 by one-way ANOVA.

To determine the relative amount of mtHA tag mitochondria in the HA immunoprecipitants, we probed for the HA tag and normalized the expression against TOMM20 for densitometry. We found that HA expression was lower in NPHS2-mtHA pulldowns than in CDH16-mtHA pulldowns (**Fig 4 A&D**). Since we loaded each MACS column with an equal amount of kidney homogenate regardless of genotype and since HA expression was normalized to TOMM20, the higher HA content in CDH16-mtHA tissue may be due to CDH16-mtHA mitochondria carrying more copies of the MITO-Tag per mitochondrion as expression of HA from CDH16-mtHA mice, with either pulldown, was approximately 3- to 6-fold higher than in NPHS2-mtHA mice when normalized to mitochondrial content. Further, we detected little or no HA-tag in the whole kidney of NPHS2-mtHA mice, demonstrating that methods for cell-type enrichment are necessary to detect podocyte mitochondria in kidney mitochondrial assays.

Next, we probed the blots for Peroxisomal Membrane Protein (PMP70), Calnexin and Lysosomal-associated membrane protein-1 (LAMP1) to detect the presence of peroxisomes, endoplasmic reticulum and lysosomes respectively. PMP70 was not significantly increased in HA pull-downs relative to TOMM22 pull-downs and whole kidney tissue homogenate in both NPHS2-mtHA and CDH16-mtHA mice **(Fig 4E)**. Calnexin, on the other hand, was approximately 3-fold lower than in whole tissue in both HA and TOMM22 pull-downs from all mice. These differences were significant, suggesting that although some endoplasmic reticulum protein is detected in the mitochondria pull-downs, it is not enriched relative to whole tissue **(Fig 4F)**. Both PMP70 and Calnexin relative expression levels were consistent between NPHS2-mtHA and CDH16-mtHA mouse samples. This was not the case for LAMP1 expression, which was the only organelle marker protein to show a significant difference between kidney cell types. Densitometry of LAMP1 normalized to TOMM20 levels from NPHS2-mtHA mice showed a greater than 6-fold increase in LAMP1 protein detected in HA-pulldowns relative to TOMM22 or whole tissue **(Fig 4G)**. Thus, podocyte-specific mitochondrial preps contain significantly higher levels of lysosomal protein than any other prep, indicating enrichment by HA-precipitation that is not detected in tubule cell mitochondria **(Fig 4G)**. Immunoblotting for cytoplasm proteins, Glyceraldehyde 3-phosphate dehydrogenase (GAPDH) and β-Actin, showed no enrichment for cytoplasm contamination in any of the pulldowns relative to whole tissue, further demonstrating specificity of the LAMP1 protein levels pulled down with podocyte mitochondria.

### Mitochondrial respiration declines in aged podocyte mitochondria from male, but not female mice

To test mitochondrial function specifically in podocytes and tubules we analyzed enriched mitochondria versus whole kidney mitochondria from the NPHS2-mtHA and CDH16-mtHA mouse lines. To test our hypothesis that mitochondrial function would decline with age, we measured respiration in isolated mitochondria from male and female mice at young adult age (5-6 mo) and late adult age (24-25 mo). To examine differences in the total mitochondria of each sample, we measured citrate synthase (CS) activity. CS is the first enzyme in the citric acid cycle, catalyzing the reaction of acetyl coenzyme A with oxaloacetate to form citric acid, and because CS is found only in the mitochondrial matrix it can be used as a direct measurement of mitochondrial content. In males there was no significant difference in CS activity for total mitochondrial content between HA-isolated mitochondria and TOMM22-isolated mitochondria from NPHS2-mtHA mice when measured in young or aged mice (**Fig 5A**). This was congruent with our immunoblots against mitochondrial proteins in male mice **(Fig 4)**. When comparing mitochondria pulldowns from young versus old, however, there was a significant increase in mitochondrial content with age in male NPHS2-mtHA mice (**Fig 5A**). In females, mitochondrial content from HA-isolated mitochondria was significantly lower than TOMM22-isolated mitochondria in both young and aged NPHS2-mtHA mice (**Fig 5B**) with the only apparent change with age being a small, but significant, increase with age in CS of TOMM22 pull-down (total kidney) mitochondria. We expected that in CDH16-mtHA mice the HA pulldown would be like that of TOMM22 pull-down since total kidney isolates likely reflect tubule mitochondria above any other cell type. Confirming our expectations, there were no changes in mitochondrial content between HA-isolated and TOMM22-isolated mitochondria from CDH16-mtHA mice in either males or females (**Fig 5 C & D**).

**Figure 5.**
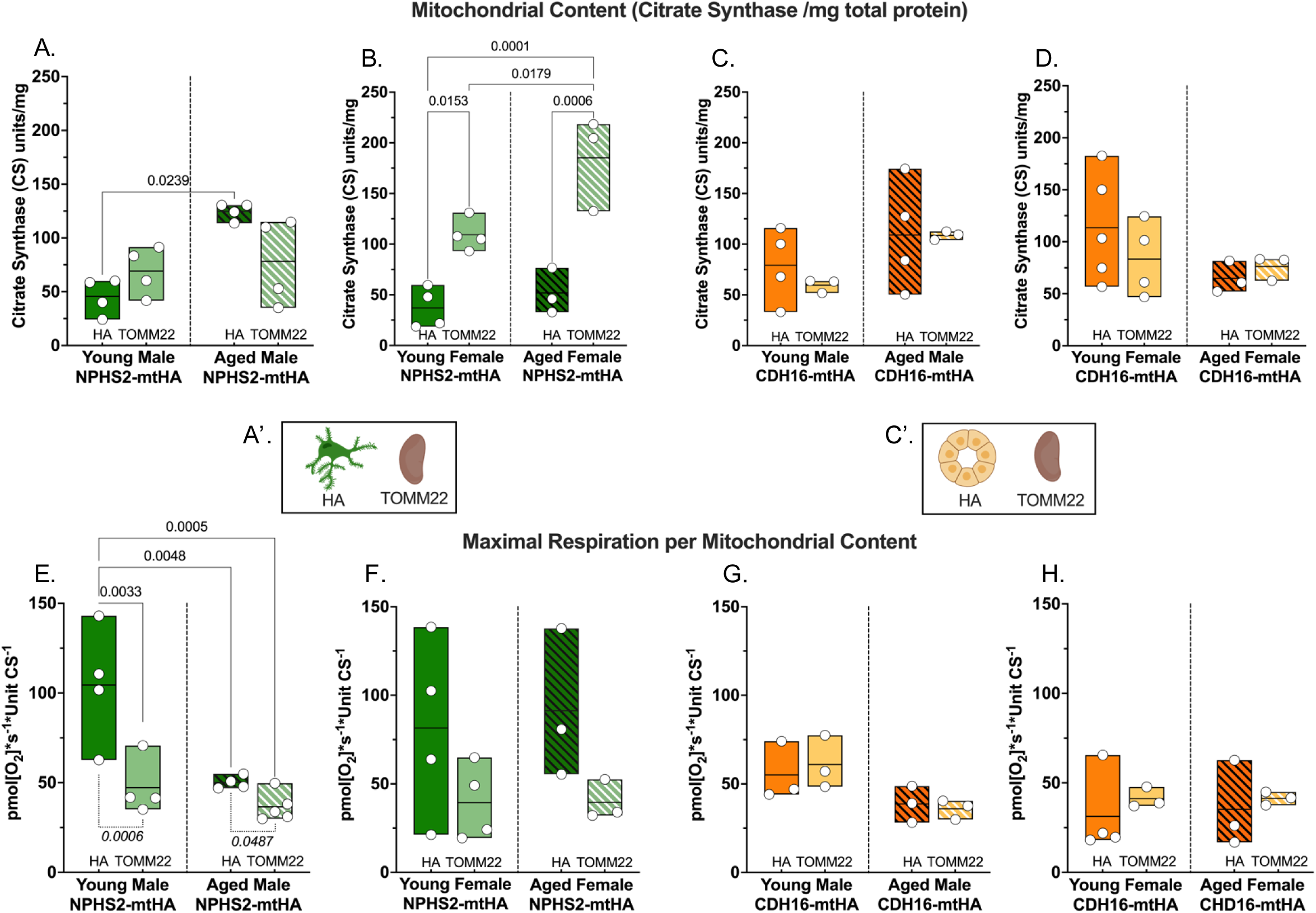
Increased respiration in podocyte mitochondria from male mice. Mitochondria isolated from NPHS2-mtHA and CDH16-mtHA mice by MACS immunoprecipitation with anti-HA (HA) or anti-TOMM22 microbeads (TOMM22) from 6-month-old and 24-month-old male and female mice were tested for respiration. **(A-D).** Total mitochondrial content measured by citrate synthase activity. No change in mitochondrial content from podocytes from male mice compared to an indiscriminate TOMM22 pulldown whereas female podocytes have decreased mitochondrial content compared to TOMM22 pulldown **(A & B)**. Significant increase in mitochondrial content with age in podocytes from males **(A)**. No differences in mitochondrial content from tubules compared to TOMM22 pulldown **(C & D). E – H.** Maximal oxidative phosphorylation (OxPhos) respiration normalized to mitochondrial content. Podocyte mitochondria from young males have significantly higher OxPhos than mitochondria using TOMM22 pulldown and OxPhos decreased with age **(E).** No differences in females **(F**) or in tubule mitochondria **(G & H)**.

We tested the enriched mitochondria for function using high-resolution respirometry for maximum state 3 oxidative phosphorylation. Based on the prevailing literature, we expected to find that respiratory capacity would be highest in tubule-specific pulldowns (CHD16-mtHA) and that these rates would be similar the whole kidney TOMM22 pulldowns, reflecting the high tubule cell content of the whole kidney. Conversely, we expected that podocyte-specific mitochondria would have a less robust respiration than tubule mitochondria when stimulated with the same substrates. Instead, we found that when normalized to mitochondrial content there was a significant increase in respiration of HA-isolated podocyte mitochondria compared to TOMM22-isolated total mitochondria from the same young male NPHS2-mtHA mice (**Fig 5E**). As we predicted, the TOMM22-isolated mitochondria from young male NPHS2-mtHA mice (47.28 ± 15.09 pmol O_2_ /second/unit CS) had a similar respiratory capacity to young male CDH16-mtHA tubule-specific (HA pulldown) mitochondrial isolations (55.04 ± 16.54 pmol O_2_ /s/unit CS) and TOMM22 mitochondria isolations (60.98 ± 14.88 pmol O_2_ /s/unit CS) **(Fig 5E &G)**. In contrast, podocyte mitochondria (HA pulldown) from male NPHS2-mtHA mice had almost twice the respiratory capacity of tubule mitochondria (104.5 ± 33.01 pmol O_2_ /s/unit CS) **(Fig 5E)**. This increased respiratory capacity in podocyte mitochondria declined with age as podocyte mitochondria from 24-26 mo aged male NPHS2-mtHA mice had half the respiratory capacity (50.01 ± 3.06 pmol O_2_ /s/unit CS) of that of young males **(Fig 5E)**. In CHD16-mtHA mice, maximal respiration of both HA and TOMM22 isolated mitochondria trended lower than in young mice, but this was not significant.

In females, although maximal respiration normalized to CS in podocyte mitochondria from NPHS2-mtHA mice trended higher than in TOMM22 isolated mitochondria, this was not significant **(Fig 5F)**. Unlike male mice, podocyte mitochondria did not show an age-related functional decline **(Fig 5F)**. As with male mice, no differences were found between ages or pulldown groups in maximal respiration from CDH16-mtHA female mice **(Fig 5H)**.

### Mitochondrial respiration declines in primary podocyte cell culture

Previous functional studies of podocyte mitochondrial respiration have relied on cultured podocyte cell lines or primary cultured podocytes obtained from isolated glomeruli.^15–17^ To compare the respiration of podocyte mitochondria directly isolated from the kidney via HA-pull down to those obtained from cultured podocytes, we performed a side-by-side comparison of the two methods (**Fig 6A**). For each test, both kidneys of an NPHS2-mtHA mouse were minced and the tissue was divided between the MACS HA pulldown protocol for respirometry performed the same day, or for the glomerular isolation protocol to culture podocytes. Isolated glomeruli were cultured under standard conditions (**Fig 6B**) with the exception that we reduced culture oxygen from room air (∼19%) to 5% in the culture incubators to better recapitulate normal kidney oxygen tension and therefore reduce the potential for oxidative stress on the cells and their mitochondria.^18^ After 7 days, podocyte mitochondria were isolated from cultured glomerular outgrowth cells using the same MACS HA pulldown protocol with respirometry performed immediately after. Paired comparison of the two podocyte mitochondria populations, direct isolate or post-culture, showed no difference in mitochondrial content by CS (**Fig. 6C**). Culturing podocytes, however, resulted in a significant decrease in isolated mitochondrial respiration (**Fig 6D**) and a significant increase in total peroxide production (**Fig 6E**), demonstrating that podocyte mitochondria exhibit detrimental functional change after only one week in primary culture.

**Figure 6.**
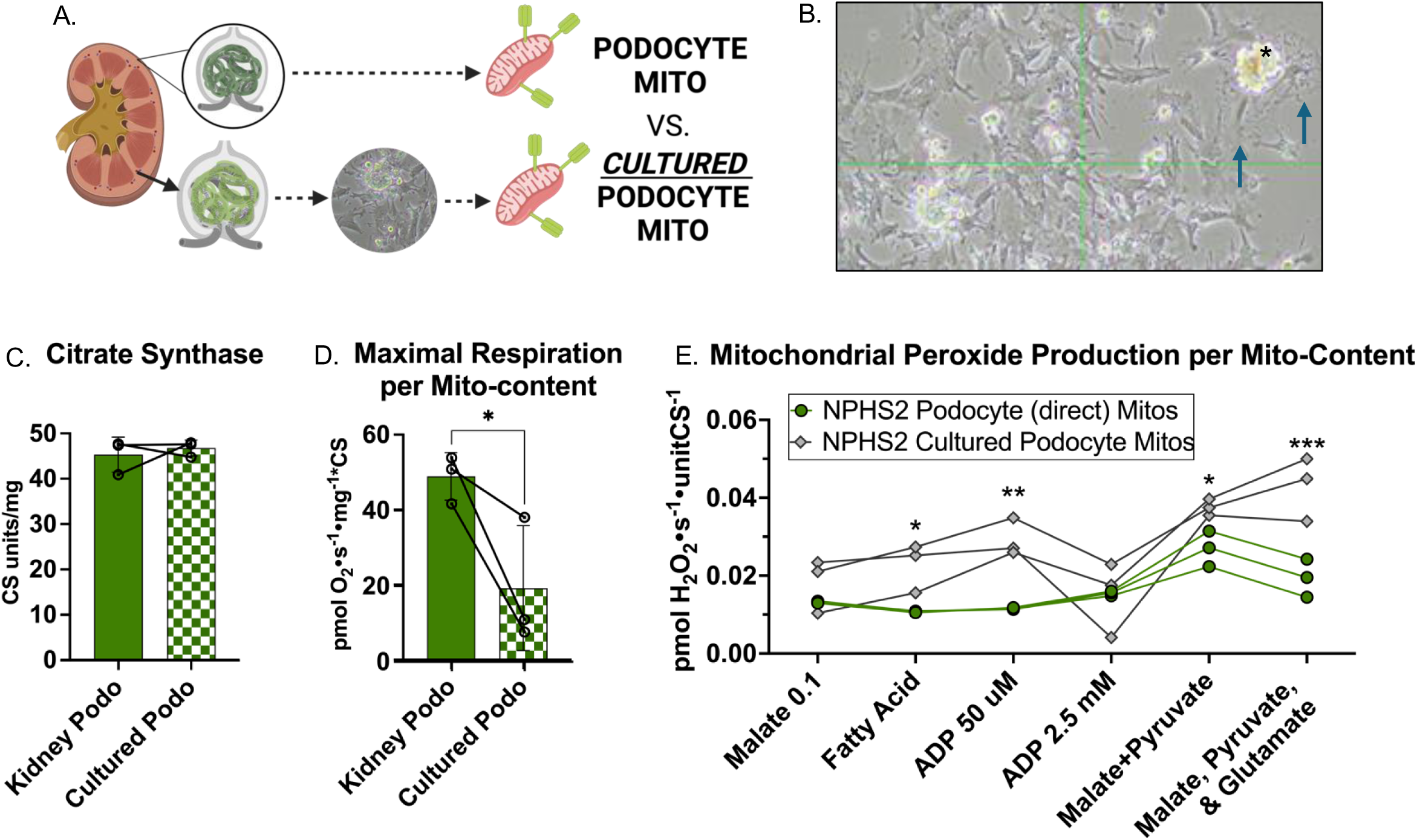
Podocyte mitochondria respiratory capacity is diminished by standard primary cell culture methods. **(A)** Experimental design of side-by-side comparison using isolation of mitochondria from either fresh tissue or cultured podocytes. The kidneys from the same animal (female, 8 mo) were used as the source of podocytes to be processed either for immediate mitochondrial respirometry from HA-tagged mitochondria or after 7 days following podocyte outgrowth from isolated glomeruli. **(B)** Podocyte growth in tissue culture conditions after glomerular isolations, arrows show examples of primary podocytes next to a glomerulus (asterisk). After one week of culturing, podocyte mitochondria were isolated following the same protocol used for whole tissue and measured for respirometry. **(C)** Mitochondrial content measured by citrate synthase activity in each preparation was not different between fresh isolate and cultured podocytes. **(D)** Maximum oxidative phosphorylation respiration normalized to mitochondrial content was significantly reduced in cultured podocytes. **(E)** Mitochondrial peroxide production was higher in mitochondria from cultured podocytes. Error bars = +/- SD, statistical significance of <0.05 determined by paired two-way t-test **(C&D)**, 2-way ANOVA with Tukey correction for **(E)**.

## DISCUSSION

These results demonstrate the importance of cell-specificity to studying mitochondria function in kidney, particularly in the podocyte, a non-proliferative cell of high importance, but low abundance, in the kidney. Our mouse experiments were designed to capitalize on the use of a promiscuous TOMM22 versus directed HA pulldown (**Fig 1**), so it naturally included an internal control for comparisons between podocytes and tubules. It is important to note that rodent models do not perfectly recapitulate human pathophysiology. In humans, chronic kidney disease (CKD) is more prevalent in women^19,20^ but men are more likely to die due to complications from CKD^21^. It has been hypothesized that this disparity may be related to higher prevalence of comorbidities due to riskier lifestyle factors among men^22,23^ and/or differences in response to diagnosis or choices for treatment following diagnosis^24–26^. However, despite these differences, multiple studies have demonstrated that renal function declines faster in males than females^27–29^. This is consistent with our results suggesting some renal pathology with age intrinsic to males may include age-related mitochondrial functional decline in podocytes that is not present in females (**Fig 5E &F**).

Interestingly, there was not an age-related decline of function in mitochondria from tubule cells (**Fig 5G &H**). This may reflect differences in proliferation capacity as well as mitochondrial dependance in tubules versus podocytes. Kidney has an absolute number of non-proliferative podocytes per glomerulus^30,31^ and both the absolute and relative number of podocytes decrease with age^5,32^. In our novel mouse crosses, functional GFP in the mitochondria of the podocyte was sufficient to cause abnormal mitochondrial structure when expressed in the podocyte, but not when expressed in the CDH16 positive tubules. The resulting phenotype was so severe as to cause early mortality (**Fig 2A**) from kidney dysfunction that was podocyte directed as indicated by increased proteinuria (**Fig 2C**) and disruption of filtration slits (**Fig 3B &C**). In this study we did not explore further the mechanism underlying how the functional GFP tag (but not the mtHA tag, which also included a non-functional GFP) was detrimental only when expressed in the podocyte. Had we not developed the mice for aging studies, and instead analyzed the mice from 6-8 weeks of age as is common practice in the field, it is possible that we would have missed the kidney dysfunction that developed in the NPHS2-mtGFP mice. Our long-term and late age studies highlight that the mtHA tagged mice are an appropriate model to measure physiologic kidney cell-specific mitochondria.

Our finding in the NPHS2-mtGFP mice also underscores the importance of cell organelle maintenance in long-lived post-mitotic cells. Other studies have pointed to mitochondrial maintenance as a priority for podocytes. Turnover of mitochondrial proteins is higher in podocytes than in the rest of the kidney^36^. It has also been shown that autophagy, in which cell debris and damaged mitochondria are engulfed by an autophagosome, before fusing to a lysosome for degradation, is higher in podocytes relative to other kidney cells^37,38^. Clinically, lysosomal regulation of podocyte health is implicated in nephropathy as with Fabry disease^39,40^ and minimal change disease^41^. Intriguingly our data also point to a stronger relationship between lysosomes and mitochondria in podocytes than in tubule cells **(Fig 4A &G)**. Although our immunoblot data contained only male mice, we recently confirmed this co-enrichment of podocyte mitochondria with lysosomes from a separate study using female cohort of our kidney MITO-Tag mice in conjunction with cross-linking proteomics.^42^

To date, common strategies for research in cell-specific bioenergetics of the kidney have been to use either cell culture models, sorted primary cells, or isolated mitochondria from whole kidney lysates from rodent models^9,15,16,43^. In a study from Brinkkoetter et al., podocyte specific knockout in mice of PGC1-α, (mito-biogenesis), Drp1(mito-fission), and Tfam (mitochondrial transcription) showed that genes once-considered essential for regulating mitochondrial functions resulted in no overt glomerular dysfunction *in vivo*^15^. This suggested that mitochondria in podocytes contributed less to energetic demand than other ATP generating systems such as glycolysis. Although the study used mice aged to at least 12 months-old and some lines to 24 months, kidneys were not stressed and measurement of mitochondrial function in podocytes was entirely reliant on podocyte disassociation and culturing. In the same study *in vitro* experiments, showed that podocyte knockdown of Tfam resulted in decreased mitochondrial respiration from primary podocytes. As well, loss of either Tfam or PGC1a in primary cultured human podocytes increased reliance on glycolysis^15^. We suspect that in these studies the act of isolating podocytes for culture acted as a ‘second hit’ stressor so that the injury to the podocyte via mitochondrial disruption was detected *in vitro*, but not *in vivo*. Our findings here demonstrate that culturing podocytes creates a baseline of mitochondrial stress (**Fig 6 D & E**). What sets our current study apart from previously published results is the use of HA-tagged mitochondria in cell-specific populations (**Fig 1**). This allowed for directed mitochondrial isolation without the need for stressful and damaging cellular dissociation. Until now it was assumed that podocytes were not heavily dependent on mitochondria for energy production, however these results demonstrate that even given the relative paucity of mitochondria in the total kidney they may in fact be much more important for meeting bioenergetic needs than previously thought. Our model revealed a significant increase in mt-HA male podocyte derived mitochondrial respiration compared to non-specific TOMM22 pulldowns (**Fig 5E**) and a trend of increased respiration in females (**Fig 5F**). Further comparison of direct isolation of mitochondria from podocytes using mt-HA pulldown revealed that respiration is twice as high when compared to commonly used primary podocyte *in vitro* culture models (**Fig 6D**).

Together these data suggest that mitochondria in the podocyte have been misclassified as unimportant for overall podocyte function and kidney health. This was likely due to the difficulty to directly examine podocyte mitochondria without the interference of the much more common mitochondria from tubules. The use of these novel mt-HA tagged models has removed that barrier, however it also raises the possibility that until now podocyte mitochondria have been under regarded for their importance in kidney health and function. Future studies of mitochondria and kidney function should focus not only on sexual dimorphism, but also include direct comparisons of podocyte to tubule mitochondrial function.

## Acknowledgements

The authors wish to thank Dr. Sarah Huen (University of Texas Southwestern) for helpful discussions and shared reagents. 3D-SIM images and quantification for filtration slit density was performed by NIPOKA GmbH. Some figures used Biorender.com for the platform to create the schematics in this manuscript. This work was funded by the National Institutes of Health: National Institute on Aging (K01AG062757 to MTS), (K01AG073470 to MSC), Nathan Shock Center (NIA P30 AA013280 to DJM); National Institute of Diabetes and Digestive and Kidney Diseases (R21DK128540 to MTS and MSC)

