## Supplemental Data for "Mitochondrial respiratory capacity in kidney podocytes is high, age-dependent, and sexually dimorphic"

| Protein of Interest | Company | Location |
|---|---|---|
| HA | Novus | Centennial, CO, USA |
| TOMM20 | Novus | Centennial, CO, USA |
| ATP5a1 | ThermoFisher | Rockford, IL, USA |
| Calnexin | ThermoFisher | Rockford, IL, USA |
| B-Actin | Cell Signaling | Danvers, MA, USA |
| GAPDH | Cell Signaling | Danvers, MA, USA |
| LAMP1 | Invitrogen | Carlsbad, CA, USA |
| PMP70 | Abcam | Cambridge, UK |

| Catalog Number | Dilution Used |
|---|---|
| NB600-362 | 1:8750 |
| NBP1-81556 | 1:7500 |
| PA5-27504 | 1:3000 |
| PA5-34754 | 1:20000 |
| 4970 | 1:1000 |
| 5174 | 1:1000 |
| 14107182 | 1:1000 |
| ab85550 | 1:1000 |
